# Accelerating the Calculation of Protein-Ligand Binding Free Energy and Residence Times using Dynamically Optimized Collective Variables

**DOI:** 10.1101/417915

**Authors:** Z. Faidon Brotzakis, Vittorio Limongelli, Michele Parrinello

## Abstract

Elucidation of the ligand/protein binding interaction is of paramount relevance in pharmacology to increase the success rate of drug design. To this end a number of computational methods have been proposed, however all of them suffer from limitations since the ligand binding/unbinding transitions to the molecular target involve many slow degrees of freedom that hamper a full characterization of the binding process. Being able to express this transition in simple and general slow degrees of freedom, would give a distinctive advantage, since it would require minimal knowledge of the system under study, while in turn it would elucidate its physics and accelerate the convergence speed of enhanced sampling methods relying on collective variables. In this study we pursuit this goal by combining for the first time Variation Approach to Conformational dynamics with Funnel-Metadynamics. In so doing, we predict for the benzamidine/trypsin system the ligand binding mode, and we accurately compute the absolute protein-ligand binding free energy and unbinding rate at unprecedented low computational cost. Finally, our simulation protocol reveals the energetics and structural details of the ligand binding mechanism and shows that water and binding pocket solvation/desolvation are the dominant slow degrees of freedom.

## 1 Introduction

Elucidating the way a drug interacts with its molecular target is of paramount relevance for drug design. The development of reliable models of ligand-protein interaction has lead to the discovery of new compounds.^1,2^ In this context, a plethora of methods aimed at identifying the ligand binding mode and estimating the protein-ligand binding free energy has been reported, see for instance recent reviews at Refs.^1,2^ From a technical point of view, we can group the available methodologies into three categories: i) molecular docking;^3–5^ ii) end-point methods such as linear interaction energy,^6^ Poisson–Boltzmann surface area,^7^ Generalized Born Surface Area; ^8^ and iii) free energy pathway methods such as Free Energy Perturbation, ^9^ Thermodynamic Integration,^10^ Transition Path Sampling,^11^ Umbrella Sampling,^12^ Steered-MD,^13^ and Funnel-Metadynamics (FM).^14^

Although docking methods are very fast in screening and ranking ligand bound poses, they suffer from inefficient sampling and scoring function inaccuracy. One of the major limitations of docking is the lack of solvation and protein flexibility dynamics. In order to obtain accurate binding free-energy, one might use either end-point methods or free energy-pathway methods. Despite the robustness of these methods, most of them require a priori knowledge of the ligand binding mode and assume that the ligand studied have similar proprieties (e.g., Free Energy Perturbation), thus hampering general applicability.

At variance with the other methods, FM does not need any preliminary information on the ligand binding mode. Once the approximate location of the binding pocket is defined, the user is able to capture the whole ligand binding mechanism from the fully solvated state to its final binding mode. The calculation of a well-characterized binding free-energy surface (BFES) allows identifying the lowest energy state (i.e., ligand binding mode) and other energetically relevant minima, which might represent important alternative binding poses. Furthermore, the absolute protein-ligand binding free energy is accurately computed from the free-energy difference between the ligand bound and unbound free energy minima. FM has been successfully used in a wide number of studies to investigate ligand binding to both proteins and DNA.^14–16^

Like in standard metadynamics, in FM a history dependent bias is added to the system Hamiltonian as a function of a set of Collective Variables (CVs), thereby enhancing the fluctuations along those CVs that mediate the rare transitions. The choice of CVs is critical. Bad CVs lead to to hysteresis and poorly converged calculations. ^17,18^ Unfortunately, the simulation time required to converge the free energy increases exponentially with the number of CVs, thus limiting the choice of the system’s degrees of freedom that can be included. This scenario poses first the need to discover the slow degrees of freedom to bias, and then develop CVs able to describe these modes. Here, we have pursued this goal by combining two enhanced sampling techniques: the Variation Approach to Conformational dynamics Metadynamics (VAC-MetaD) ^19^ with the Funnel-Metadynamics. The first, allows capturing the details of the intermolecular interactions during binding while the second ensures an exhaustive exploration of the bound and unbound state and a faster convergence.

The benzamidine/trypsin system has been widely studied with different computational techniques.^14,20–23^ Its solvent exposed binding site makes it an ideal system for testing new ligand-protein binding methods. Therefore we choose benzamidine-trypsin complex as case study ^14,20,23^ using a basis set that includes general descriptors of the protein-ligand binding process. The combination of the two methods allows describing the binding mechanism of benzamidine to trypsin. We demonstrate the ability of FM using VAC-MetaD optimized CVs to reproduce the known ligand binding mode and to provide an accurate estimation of the absolute protein-ligand binding free energy, reducing the convergence time. Furthermore, this simulation provides a high resolution free-energy surface that highlights the key role played by the water molecules in the different energy minima visited by the ligand during its binding path. We also show that using these optimized CVs with infrequent deposition of the bias as reported in,^23^ leads to an accurate estimate of the ligand unbinding rate koff. The flexibility of the method permits to include in one CV a number of molecular descriptors of the protein-ligand binding process such as the number of protein-ligand contacts, ligand solvation, pocket solvation and ligand orientation. The analysis of the results indicates which degrees of freedom play a major role during ligand binding. In our case, we have found that the slowest degrees of freedom are the desolvation of the binding site and of the ligand rather than the formation of direct protein-ligand contacts.

## 2 Methods

### 2.1 Metadynamics

Metadynamics (MetaD) is an established enhanced sampling method used to study rare events.^17,24,25^ MetaD accelerates the sampling by adding a history-dependent bias in the form of Gaussian kernels on selected system’s degrees of freedom, also known as collective variables (CVs).

In well tempered Metadynamics ^26^ this scope is achieved by periodically adding a bias that is updated according to the iterative procedure

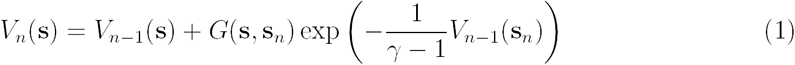

where V_*n*_(s) is the total bias deposited at iteration *n* and is obtained by adding at the previous bias V_*n-*1_ a contribution that results from the product between a Gaussian kernel *G*(s, s_*n*_) and a multiplicative factor exp *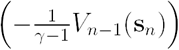* that makes the height of the added Gaussian diminished with time. The bias factor *γ* > 1 determines the rate with which the added bias decreases and regulates the amplitude of the s fluctuations. Thanks to the MetaD potential, the system explores the FES escaping the free-energy minima and crossing large energy barriers in a reasonable computational time.

An additional advantage of well tempered MetaD is that after a transient period the simulation reaches a quasi-stationary limit in which one can estimate the expectation value of any observable ⟨*O*(**R**) ⟩,^27^ as an average over the simulation time according to:

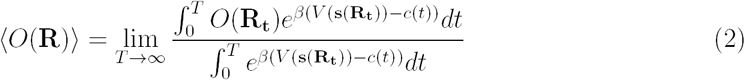

Here, **R**_*t*_ are atomic positions at time *t*, *β* is the inverse of the temperature multiplied by the Boltzmann constant *k*_*B*_ and the time dependent offset *c(t)* is defined as:

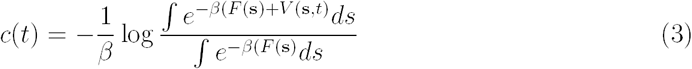

where F(s) is the free energy as a function of the CV. Following this procedure, one can reweight to Boltzmann averages of any variable even if they are not included in the biased CV.

### 2.2 Funnel-Metadynamics

The binding process between a ligand and its molecular target is typically represented as chemical reaction with the equilibrium constant *K*_*b*_ (Fig. 1A). The latter is the ratio between the concentration of the system in the bound state and that in the unbound state. In thermodynamics, *K*_*b*_ is related to the the absolute protein-ligand binding free energy *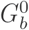*, which is the free-energy difference between the unbound and the bound state, through the formula:

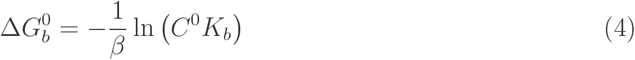

**Figure 1:**
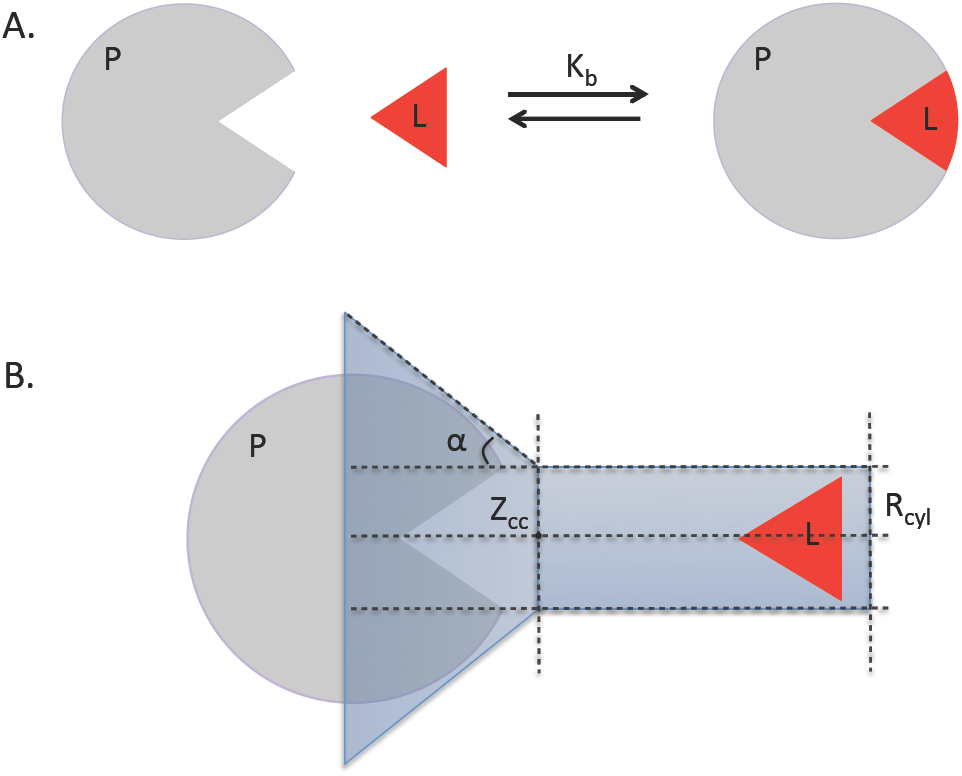
A) A cartoon representation of a ligand (L) binding to a protein (P). B) The funnel restraining potential illustrated in blue, along with its parameters. More specifically Z is the exit binding path axis, Z_*cc*_ is the distance along the axis Z, where the cone restraining potential turns into a cylindrical shape, and R_*cyl*_ is the radius of the cylinder, and *α* is the angle defining the cones amplitude.

where *C* ^0^=1/1660 *Å*^−3^ is the standard concentration. However, calculating the free energy along a binding transition is difficult. In particular, in the case of protein/ligand binding an additional complication arises from the fact that once the ligand leaves the protein, it has to explore a large conformational space. A remedy to this problem has been suggested in Refs.^14,28,29^ with the imposition of a restraint potential in the binding site. In this study, we will use the Funnel-Metadynamics method, where the restraining potential has in the shape of a funnel as illustrated in Fig. 1B. The design of the funnel is such that the restraint does not act on the ligand when in the binding site while confines the ligand to a cylindrical region once outside. One can also look at this procedure as having imposed an entropic restraint. The net effect of the cylindrical restraint can be accounted for, leading to the following formula:

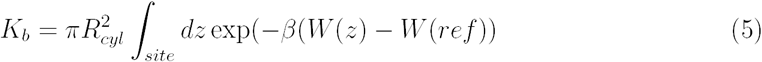

where *W(z)* and its value in the unbound state *W(ref)* can be derived from the Potential of Mean Force (PMF) along the funnel axis *Z*, while *R*_*cyl*_ is the radius of the cylindrical part of the funnel potential and *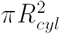* accounts for the restraining potential in the unbound state (see Ref.^14^ for details). Finally, one can calculate the absolute protein-ligand binding free energy 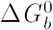 combining Eq. 4 and Eq. 5. Due to the funnel restraint, several forth-and-back events between the bound and the unbound state are observed during the simulation leading to the free-energy convergence of the calculation in a shorter computational time. ^14,15,30–33^

### 2.3 Variational Approach to Conformational dynamics Metady-namics

As already discussed, in MetaD calculations the number of CVs should be kept as small as possible, since the cost of reconstructing the free energy grows exponentially with the number of CVs used. This limitation is particularly severe in protein systems where a slow process often involves several degrees of freedom. Recently Mc Carty and Parrinello developed a Variational Approach to Conformational dynamics in Metadynamics (VAC-MetaD) ^19^ that allows combining several degrees of freedom into a number of CVs that are a combination of many descriptors. So far this has lead to a better understanding of chemical reactions and folding of small peptides. ^34,35^

Here we briefly review the steps needed to apply this procedure. First one identifies a set of N descriptors *d*_*k*_(**R**) that one thinks are important for the phenomenon under investigation. Then one searches for the best CV s_*i*_(**R**) that can be expressed as a weighted linear combination of *d* _*k*_(**R**), with corresponding coefficients *b*_*ik*_:

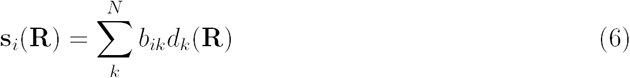

This is obtained by a variant of Time-lagged Independent Conformational Analysis (TICA) developed in Noe’s group. ^36,37^ In this setting, one computes the time-lagged covariance matrix *C*(*τ*) defined as:

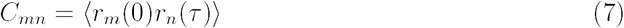

where *r* _*k*_(*τ*)=*d* _*k*_(*τ*)-⟨*d*_*k*_(*τ*) ⟩. We then determine the *b*_*ik*_ coefficients by solving the generalized eigenvalue equation:

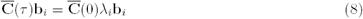

where *λ*_*i*_ is the *i*th eigenvalues, **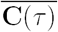** is the time lagged covariance matrix, 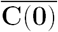 is the covariance matrix at time zero. In the variant of TICA proposed by Noe’s group., ^36,37^ they assume a long trajectory which spontaneously transits between metastable states. Mc Carty and Parrinello^19^ noticed that a similar procedure can be applied from a MetaD trajectory, provided that the MetaD timescale is properly rescaled, as follows:

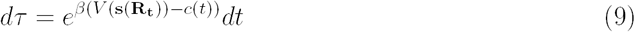

For the purpose of studying a ligand unbinding transition, the eigenvectors of the eigenvalue equation obtained from a MetaD trajectory, correspond to the slowest decaying modes, and can be used as CVs in the production MetaD simulation.^19,38^

### 2.4 Ligand unbinding rate k_*off*_ from Infrequent Metadynamics

Inspired by Voter^39^ and Grubmüller,^40^ in a recent study Tiwary and Parrinello^41^ introduced the Infrequent Metadynamics (InMetaD) framework illustrating how MetaD could be used to calculate transition rates of activated processes. The assumptions of this method are that i) there is no bias deposited in the transition state ensemble (TSE) region and ii) that the biased CVs are able to distinguish between relevant metastable states. This goal can be achieved by using an infrequent bias deposition which reduces the probability to add bias to the short transition time where the system is in the TSE region. ^41^ Metadynamics accelerates the dynamics by a factor *α*, given by a running average obtained through MetaD as:

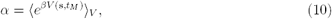

where *t*_*M*_ is the Metadynamics escape time, s is the biased collective variable at time *t*_*M*_, *β* is the inverse of *k* _*B*_T and *V(s,t*_*M*_ ) is the bias deposited at time *t*_*M*_. The real time *t* can be related to *t*_*M*_ via Eq. 11.

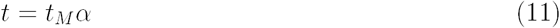

A rigorous quantitative way to assess the method’s assumption is to perform a Kolmogorov-Smirnov statistical test. ^42,43^ This effectively measures if the *M* observed escape times follow a Poisson distribution, as expected in a rare event scenario.

### 2.5 Protocol

In this section we present a schematic description of our protocol. The protein and water are modeled using the Amber-14SB and TIP3P classical forcefields, while the ligand using the Amber GAFF library.^44,45^ At variance with our previous studies, ^14,23^ here we use a new benzamidine force-field parametrization based on quantum mechanics calculations. These calculations show that in the ground state of the isolated benzamidine the amidine group has a non planar torsion angle with respect to the benzene ring. Similar results have been found in a previous study by Pophristic et al ^46^ and through X-ray diffraction by Liebschner et al.^47^ All of our simulations start from a solvated and equilibrated configuration of the benzamidine-trypsin X-ray crystral structure (PDB ID: 2oxs). Initially, we define a simple CV, namely the distance between the ligand C7 carbon and the D189 C*γ* of the pocket. Then, we run a 500 ns FM simulation which enhances the fluctuations of the binding transition. Using this FM trajectory we perform the VAC-MetaD optimization procedure implemented in the PLUMED software^48^ to generate the CVs biased in a 280 ns VAC-FM production run. Both simulations are performed at room temperature and atmospheric pressure. For more information about the system preparation we refer the interested reader to the SI. For the sake of clarity, we describe below the VAC-MetaD procedure in the following steps.

Made wise by the many studies showing that direct ligand/protein contacts, as well as the solvation and the orientation of the ligand are involved in the stability of the intermediate states,^14,18,23,49^ we define a basis set of descriptors for ligand/protein binding (*N*_*pr/lig*_,*W*_*pock*_,*αβ*(*θ*),*W*_*lig*_). In particular, *N*_*pr/lig*_ corresponds to the protein-ligand contacts, *W*_*pock*_, corresponds to the number of water molecules in the pocket, *αβ*(*θ*) is a transformation of the torsion angle describing the rotation of the ligand relative to the protein binding site, and *W*_*lig*_ is the number of contacts between water molecules and the ligand. For more details see Table S1. After defining the basis set we perform a VAC-MetaD optimization on the first 120 ns of the preliminary FM simulation to identify the eigenvectors and eigenvalues associated with the slowest modes of the system.

We select the first two eigenvectors associated with the slowest decaying eigenvalues (see Fig. 2a,b), and use them as CVs. Since reaching convergence using one CV proves difficult (see Fig. S1) it does seem natural then to include the second eigenvector as CV, given the fact that its associated eigenvalue time decay, while faster than of the first one’s is slower than the others. Furthermore, the eigenvalues of the first two eigenvectors decay more slowly than of the other (see Fig. 2a). The two optimal CVs obtained from the VAC-MetaD analysis described above are, **s1**=−0.38 *N*_*pr/lig*_+ 0.78*W*_*pock*_ −0.19*αβ*(*θ*) +0.45 *W*_*lig*_ and **s2**=0.01*N*_*pr/lig*_ −0.85*W*_*pock*_ −0.30*αβ*(*θ*) +0.44 *W*_*lig*_. The most striking feature of this two CVs is the dominant role played by water both in pocket hydration dehydration and in ligand solvation/desolvation.

**Figure 2:**
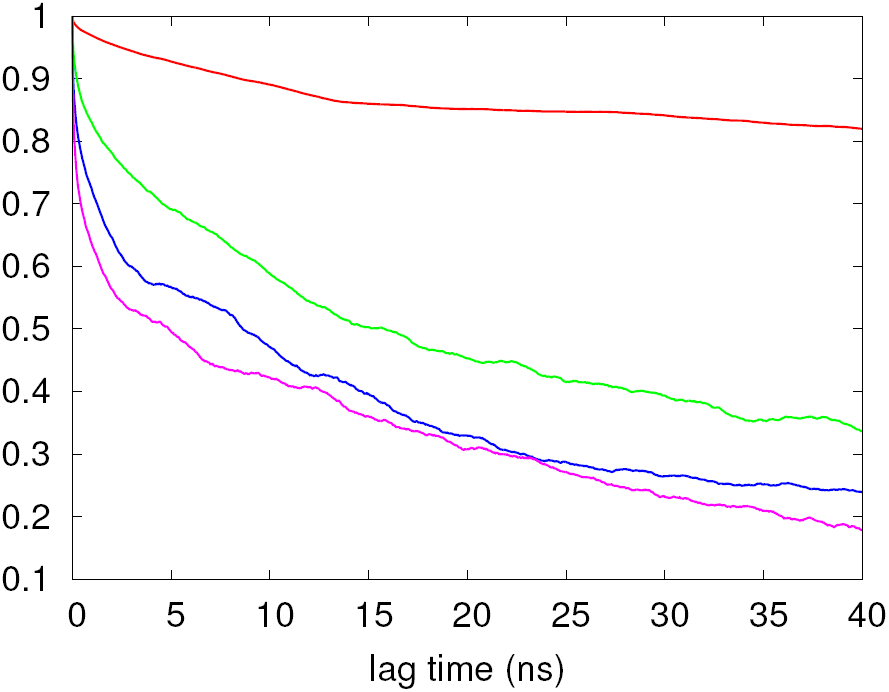
a) Eigenvalues decay as a function of lag time and b) coefficients of the basis set for the two eigenvectors associated with the two first slowest decaying eigenvalues (red and green respectively).

## 3 Results

In the following paragraphs we address the binding mechanism of benzamidine/trypsin complex, the convergence speed obtained using FM with VAC-MetaD optimized CV and finally the unbinding rate.

### 3.1 Free energy

First, we perform a production run of 280 ns FM simulation, biasing two CVs **s1** and **s2** obtained earlier. Using these two CVs we computed the corresponding free energy surface (see Fig. 3a). Three stable bound states are found, A, P and B. The first corresponds to the crystal structure, the second to a presolvated state, the third to a rotated presolvated still bound state. A detailed description of the stable bound states follows.

**Figure 3:**
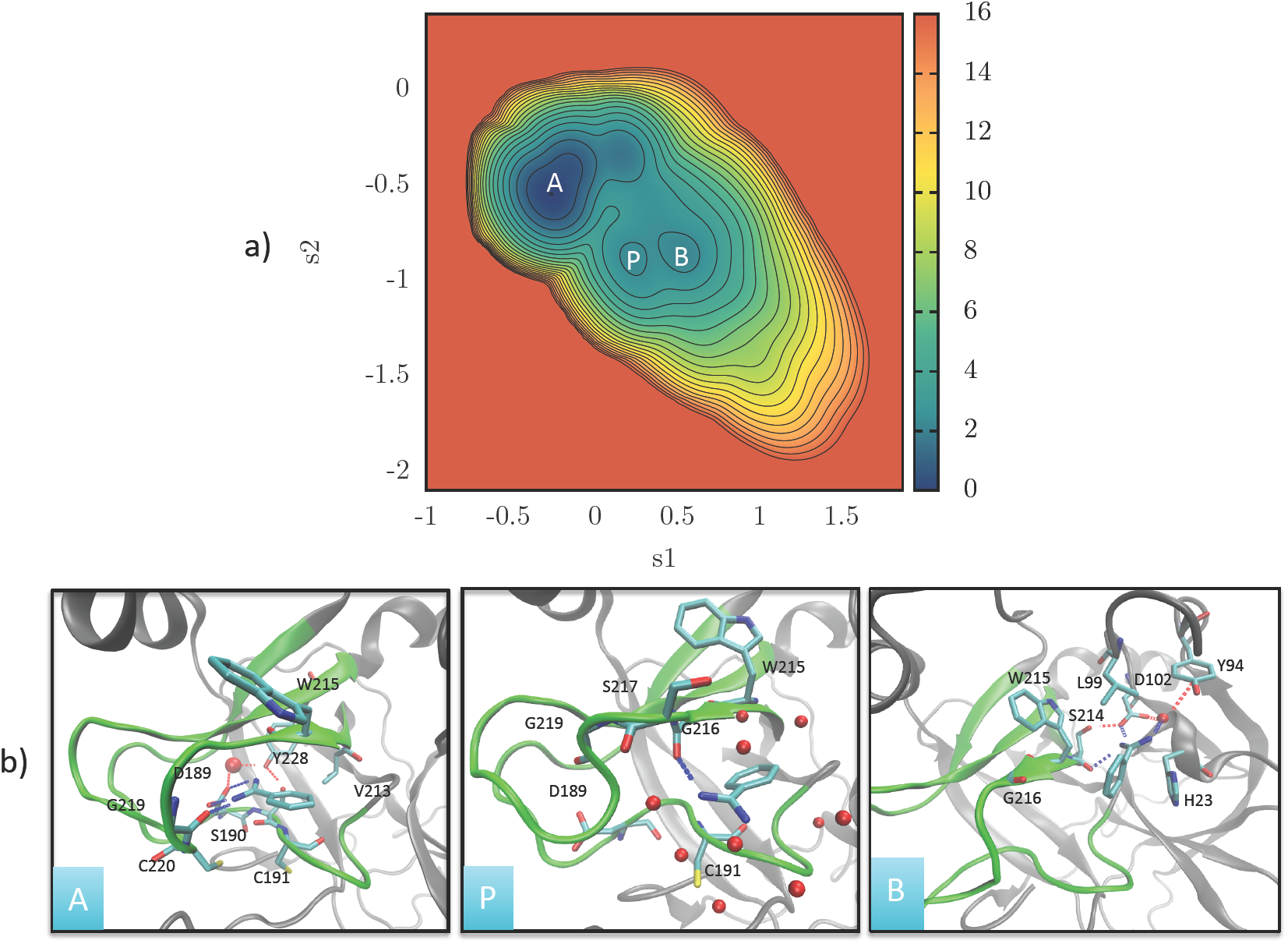
a) Free energy surface as a function of the two biased eigenvectors **s1** and **s2**. where in white the stable states and b) a detailed description of the bound poses.

#### Minimum A

Minimum A corresponds to the crystal structure. In this minimum the ligand’s diamino group forms direct hydrogen bonds with the hydrogen bond accepting groups of the negatively charged D189, polar S190 and G219. Moreover, this minimum is stabilized by a hydrogen bond network formed by a structural water, bridging the interactions of Y228, D189 and S190. Finally, the ligand’s aromatic ring, forms hydrophobic contacts with V213, C191 and W215. These interactions are in agreement with previous studies. ^14,20,23,50,51^

#### Minimum P

This minimum corresponds to a presolvated state. In P, the ligand is rotated compared to state A, with its diamino group exposed to solution. The phenyl ring of the ligand is sandwiched between the C*α* of W215 and C191. It forms direct hydrogen bond with G216 while at the vicinity of the triad G216, S217, G219. Minimum P has been also discussed in Ref.,^23^ where is also called P, and it is the same as state P2 of Ref. ^50^

#### Minimum B

This minimum corresponds to a rotated presolvated bound state. In B, the ligand is rotated compared to minimum A and sits just outside the pocket. The ligand’s diamino group forms direct H-bond with the negatively charged D102, and S214. The ligand is sandwiched between W215 and H23 forming *π*-stacking interactions.^52^ Finally, a water forms a hydrogen bond network between the diamino group of the ligand, Y94 and D102. This mimimum, is very similar to state P12 and/or TS2 in ref. ^50^ The combination of these interactions is allowed only by a rotated torsion, and can be captured by the new parametrization of the ligand.

#### Binding Free Energy Surface (BFES)

The representation of the BFES shown in Fig. 3 does not clearly describe the position of these states relatively to the unbound state region. Thus we represent the free energy landscape as a function of the same two variables used in our previous study.^14^ One, *Z*, gives the ligand’s center of mass position projected along the funnel axis. The other, Θ, is the orientation of the ligand relative to the binding site (see Table S1). In this BFES representation (Fig. 4a), A, B and P can also be clearly identified. Here, it is possible to distinguish two additional local minima A and AP in which most of the ligand/protein interactions are mediated by water molecules. These minima represent transient and preliminary binding poses that easily evolve to the energetically most stable states A and P within 20 ns of unbiased MD calculations. The interested reader can find more information about these minima in the SI.

**Figure 4:**
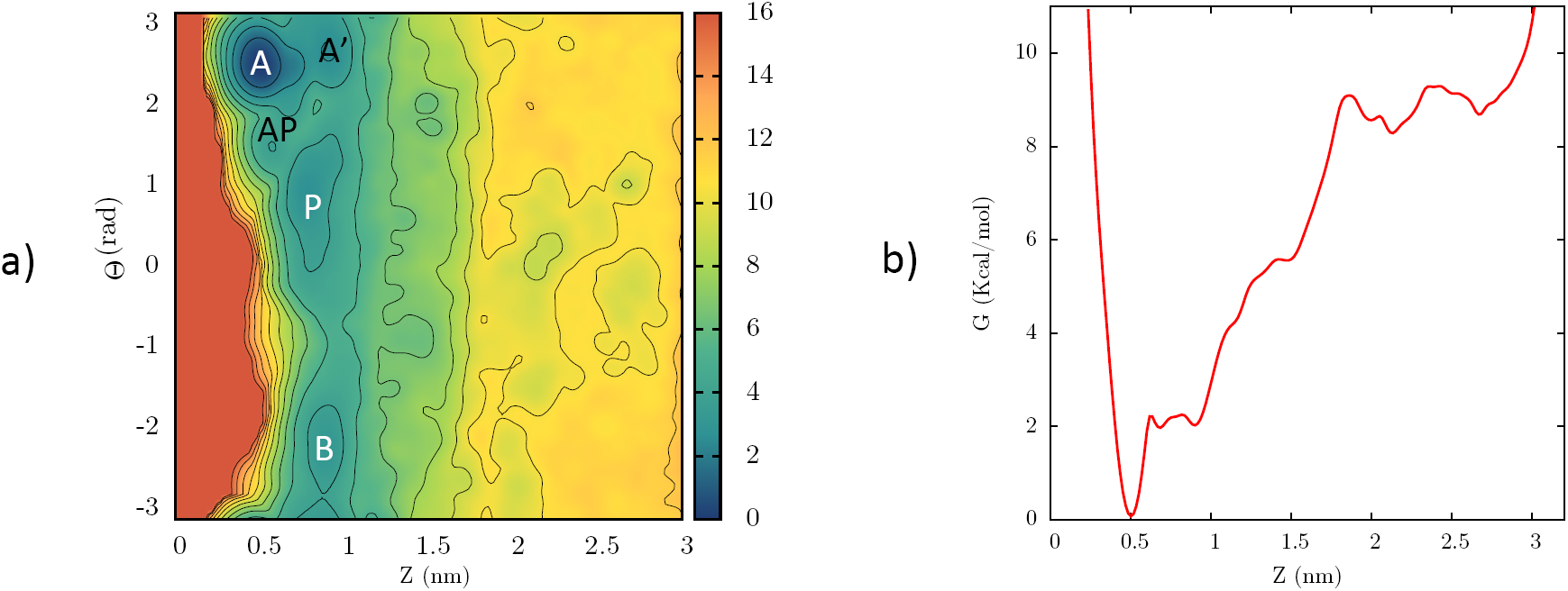
BFES as a function of the position of a) the ligand’s center of mass along the Z axis of the funnel and the torsion Θ, where in white the stable bound states, in black the unstable binding poses. b) The *Z* axis obtained from the VAC-MetaD optimized CVs FM simulation.

#### The role of water

It is known that explicit protein/water/ligand H-bond bridging interactions can increase the stability of the bound pose by several kcals/mol. ^18^ In the benzamidine system it is known ^14^ and verified here that state A is stabilized by a hydrogen bond network formed by a structural water bridging the interactions of Y228, D189 and S190. Given such a positively charged pocket, ^53^ it makes sense that upon breakage of some protein-ligand interactions of minimum A, they will be replaced by water mediated H-bonds between the ligand and the protein, thus forming nearby minima A’ and AP (see Fig. S3). Here, the stability of state P reflects this balance of solvation and direct protein-ligand forces. Importantly, at all stages of binding, water is involved in the process, be it from a ligand desolvation point of view, or from a structural water inside the pocket point of view.

### 3.2 Fast prediction of the binding fee energy

From the projection of the BFES on the *Z* axis of the funnel (see Fig. 4b) we can compute the absolute protein-ligand binding free energy 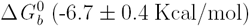, whose value after the correction in Eq. 5 gives the value in embarrassing agreement with the experimentally reported value of 6.4 to 7.3 kcal/mol. ^54,55^ It must be also underlined that the VAC-MetaD optimized CVs simulation converges smoothly to this value, while the FM with suboptimal CVs after 500 ns still exhibits relative large fluctuation (see Fig. S2). Regarding the technical aspects of this calculation we refer the interested reader to the SI.

Finally, a remarkable finding is that the the main features of the BFES as the local minima of the bound state, are already captured from the first 60 ns of the VAC-MetaD optimized CVs FM simulation (see Fig. S8d), posing a great improvement compared to the FM with suboptimal CVs where one would have to simulate at least 240 ns (see Fig. S8b) to obtain a 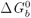 value in the right ball park.

### 3.3 Unbinding rate

Using 20 simulations initiated in state A, we perform Metadynamics with infrequent deposition of the bias (see SI) and stop the simulation once the trajectories reach the unbound state (*Z* > 2 nm). The associated *k* _*off*_ found is 4176 ± 324 *s*^−1^, which given the uncertainties intrinsic to these type of calculations, can be considered as a good agreement with the experiment *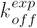* of 600 *±* 300.^56^ Note that the p-value of the KS test performed was 0.77, indicating the quality of the optimized CVs (see Fig. S9). The different result compared to Ref.^23^ can be explained by the new ligands torsion potential used in this study, which allows a much easier rotation of the ligands diamino group. In so doing, the benzamidine can disentangle itself from the interactions formed at the lowest minimum A and reach the solvated state more rapidly.

## 4 Discussion

In this study we combined for the first time Variation Approach to Conformational dynamics with Funnel-Metadynamics, in order to accurately predict the binding energy, binding mechanism, and unbinding rate of the benzamidine-trypsin complex. In this way we find ligand and pocket solvation/desolvation to be the slow degrees of freedom by looking at the slowly decaying eigenvectors of the VAC-MetaD, which when biased significantly accelerate the convergence of the bound state and the overall BFES, compared to FM with suboptimal CVs. The estimated absolute binding free energy of FM using optimized CVs is in excellent agreement with experiments. In addition it gives a better resolved BFES-including more resolved water rich local minima A’ and AP-compared to the FM simulation of suboptimal CVs. Moreover, when VAC-MetaD optimized CVs are used with the Infrequent Metadynamics scheme, it predicts the unbinding rate in good agreement with the experiment, thus providing a relatively cheap, rigorous and accurate calculation with respect to already existing methods.^56^

## 5 Acknowledgements

Z. F. B. and M. P. acknowledge VARMET European Union Grant ERC-2014-ADG-670227 for funding. V. L. acknowledges the support from the Swiss National Science Foundation (Project N. 200021 163281) and the COST action CA15135 (Multi-target paradigm for innovative ligand identification in the drug discovery process MuTaLig). All the authors acknowledge the Swiss National Supercomputing Centre (CSS) for providing the computational resources.

